# Topoisomerase IIIβ protects from tumorigenesis and immune dysregulation

**DOI:** 10.1101/2025.03.13.643061

**Authors:** Rasel Al Mahmud, Simone Andrea Baechler, Anjali Dhall, Sourav Saha, Hongliang Zhang, Shuling Zhang, Min-Jung Lee, Nahoko Sato, Shraddha Rastogi, Suresh Kumar, Muhammad S. Alam, Liton Kumar Saha, Beverly A. Mock, Valentina M Factor, Yves Pommier

**Affiliations:** Developmental Therapeutics Branch & Laboratory of Molecular Pharmacology, Center for Cancer Research, National Cancer Institute, NIH, Bethesda, MD 20892, USA; Laboratory of Cancer Biology and Genetics, National Cancer Institute, National Institutes of Health, Bethesda, Maryland, USA; Clinical Translation Unit, Laboratory of Molecular Biology, Center for Cancer Research, National Cancer Institute, National Institutes of Health, Bethesda, MD 20892, USA

**Author notes:** Equal contributors (Rasel Al Mahmud, Simone Andrea Baechler). **Author Contributions:** Y.P. supervised the study. M. R. A. M. and Y.P. conceived, managed, and oversaw the overall project, Y. P. and B. A. M. provide resources, M. R. A. M., S. A. B. and V.M.F. designed the experiments. M. R. A. M., S. A. B., V.M.F., S.S., H.Z., S.Z., M.L., N.S., S.R., S.K., M.S.A. performed research, M. R. A. M. and A. D. performed data analysis, M. R. A. M. wrote the original draft, M. R. A. M., Y.P., S.A.B., V.M.F., S. S., L.K.S reviewed and edited the manuscript, and M. R. A. M., V. M. F. and Y.P. critically reviewed and finalized the manuscript. **Declaration of interests:** The authors declare no competing interests.

**Keywords:** Topoisomerase, R-loop, DNA breaks, Immune dysregulation, Lymphoma

## Abstract

Topoisomerase III-beta (Top3b) reduces nucleic acid torsional stress and intertwining generated during RNA and DNA metabolism while protecting the genome from pathological R-loops, which otherwise result in DNA breakage and genome instability. By studying Top3b knockout mice (Top3b-KO), we find that the loss of Top3b accelerates the development of spontaneous lymphoid tumors arising in spleens and lymph nodes, the organs with prominent Top3b expression. Aging Top3b-KO mice also display splenomegaly and systemic immune alterations including neutrophilia and lymphopenia suggestive of chronic inflammation. At the molecular level, Top3b deficiency causes genome-wide R-loop accumulation in splenocytes as measured by CUT&Tag sequencing. Increased R-loops is associated with genomic DNA breaks and activation of immune signaling pathways including the IL-6 signaling, interleukin-7 signaling and cGAS-STING. Moreover, knocking-out Top3b promotes the rapid development of syngeneic EL4 T-cell lymphomas. In conclusion, our work implies that, in addition to its role in preserving the nervous system, Top3b protects from tumorigenesis and immune dysregulations.

**Significance:** Topoisomerase III-beta (TOP3B) is the only dual RNA-DNA topoisomerase. Its inactivation has previously been shown to be critical for neurodevelopmental disorders in humans and R-loop suppression in cell line models. Here, we demonstrate that loss of Top3b accelerates spontaneous lymphomagenesis and promotes syngeneic lymphoma growth in association with innate immune defects and R-loop accumulation in Top3b-KO mice. These findings provide evidence for a previously unappreciated role of Top3b as a tumor suppressor gene and immune regulator.

## Iintroduction

Topoisomerases are pivotal for preventing and resolving the nucleic acid topological constraints arising from fundamental molecular processes including DNA replication, transcription, translation, meiotic recombination and chromosomal segregation (1-4). Among the six metazoan topoisomerases (Top1, Top1mt, Top2a, Top2b, Top3a and Top3b), only topoisomerase 3 beta (Top3b) can resolve RNA topological problems, in addition to its activity on single-stranded DNA (3, 5).

Eukaryotic Top3b works as a heterotrimeric complex with the scaffolding protein Tudor domain-containing protein 3 (TDRD3) and Fragile X Mental Retardation Protein (FMRP) (6, 7). Top3b is endowed with a wide range of functions facilitating nucleic acids metabolic processes (5-13). Top3b plays a prominent role in suppressing cellular R-loops (3, 13-15) either through relaxation of hyper negative supercoils or by resolving R-loops (8) in coordination with the helicase DDX5 (3). Top3b’s auxiliary factor TDRD3, via its’ ‘Tudor domain’, can also recognize asymmetric di-methylated arginine marks in core histones (H3R17me2a and H4R3me2a) and RNA POL2 (R1810me2a), targeting TOP3B to CpG island containing active promoters to resolve R-loops (13). Top3b-mediated suppression of R-loops regulates transcription in cancer cells and neurons (13, 16). In addition, Top3b accelerates translation of a group of RNAs both in cancer cells and neurons (5-7, 9, 12, 17). Top3b is also associated with the stabilization of RNAs in human colorectal cancer cell HCT116 independently of its catalytic activity (9).

R-loops are DNA-RNA hybrid structures with a displaced DNA single-strand and variable RNA-DNA base pairs ranging between 300-2500 base pairs on average (3, 18-20). Moreover R-loops are highly dynamic in nature with an average limited half-life of 10-15 minutes (18, 21, 22). R-loops are essential for maintaining important physiological processes, including class-switch recombination (23), mitochondrial replication (24, 25), protection against promoter methylation (26-28), transcription termination (29), chromatin remodeling and chromosome segregation (30, 31), double-strand break (DSB) repair (32, 33), DNA repair in transcribed regions,(28), telomere repair (34), and centromeric functions (28, 35). However, unscheduled and persistent R-loops can lead to a range of molecular defects, including replication stress, replication fork collapse, stalling of transcription forks, replication-transcription conflicts, DNA-double-strand breaks (DSBs), genome instability (3). As a result, R-loops are associated with a number of diseases such as amyotrophic lateral sclerosis type4 (ALS4), fragile X syndrome, autoimmune diseases, and cancer (3).

Studies with Top3b-KO mice have begun to reveal the importance of TOP3B for human health. Top3b inactivation has been associated with neurological defects, including increased anxiety, reduced spatial leaning, defective synaptic transmission and deficient adult neurogenesis. Top3b-KO mice also have a shorter life-span (36-38), infertility after successive generation of homozygous pairs breeding (39), transcriptional and behavioral impairment (4, 10), accumulation of chromosomal aberrations, and are prone to autoimmune disease (40). However, little is known about R-loops, immune-and cancer-related phenotypes of Top3b-KO mice. Here, we report our exploration of novel features of the Top3b-KO mouse phenotype by combining a combination of morphological, histological, immunohistochemical, molecular and sequencing approaches that demonstrate that unresolved and/or excessive R-loops are associated with immune dysregulation and increased lymphoma development.

## Results

### Loss of Top3b accelerates spontaneous lymphomagenesis in mice

The Top3b-KO mouse line (36) (a gift from Dr. Albert C Shaw, Yale University) was rederived before introducing into the NCI animal facility. Consistent with previous report, our Top3b-KO mice showed reduced lifespan (Fig. S1*A*) and increased incidence of dermatitis (Fig. S1 *B*) (36). Because lymphoma is the most common causal factor contributing to mortality of aging C57BL/6 mice (41), we investigated the impact of Tob3b loss on the prevalence of lymphoma in two different age groups (Fig. 1*A*). At 11-15 months of age, histological analysis of spleens detected early lymphomas in 3 out of 13 Top3b-KO mice in comparison with none in the age-matched WT littermates (Fig. 1*A*). Histology and immunohistochemical (IHC) staining using cell-type specific antibodies showed expansion of B and T cells in the Top3b-KO spleens causing disruption of the follicular structures and tissue architecture typical for lymphoid disorders (Fig. 1*B*). Early splenic lymphomas were characterized by atypical lymphoid hyperplasia of pale, slightly enlarged immature lymphocytes causing enlargement and a partial fusion of white pulp areas (Fig. 1*C*).

**Fig. 1.**
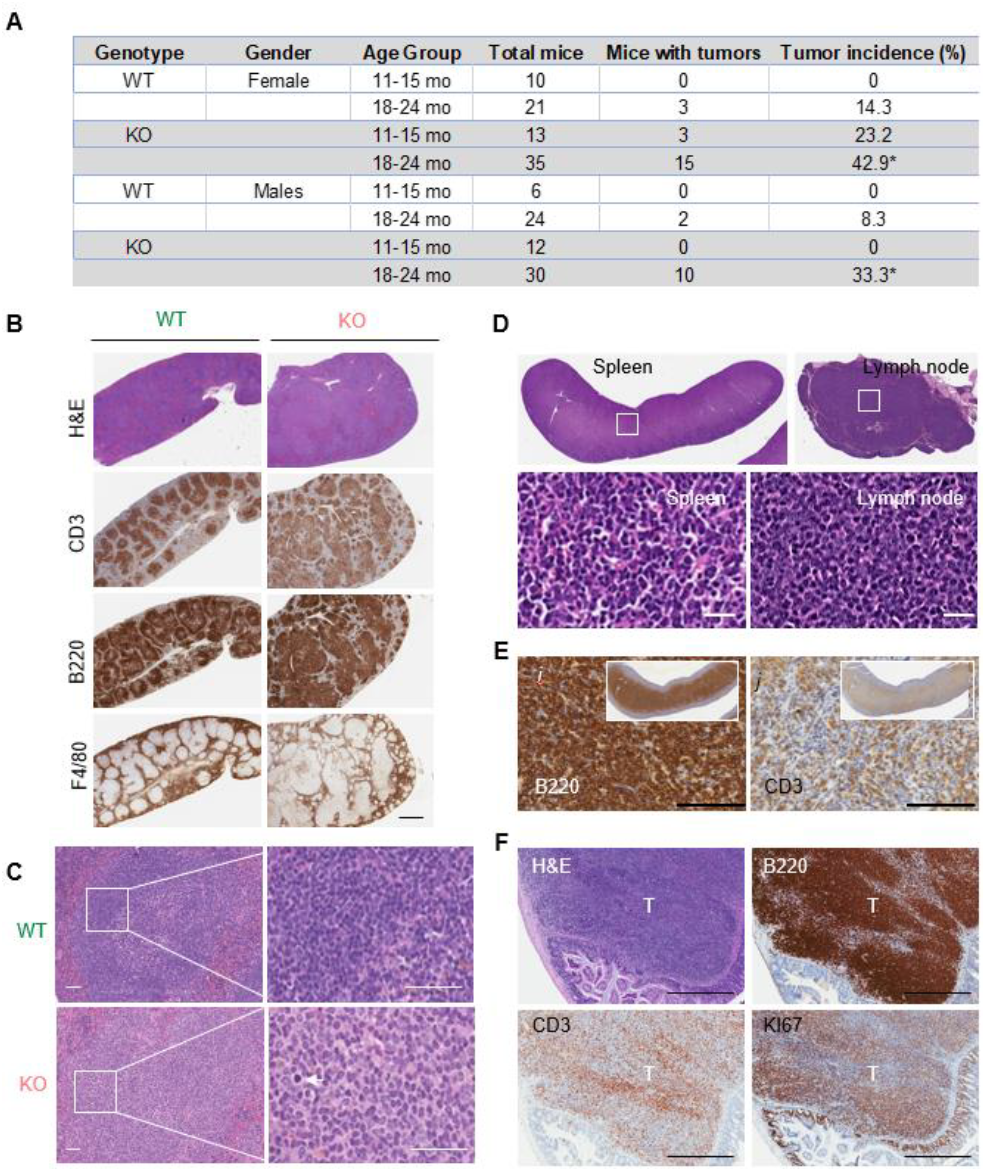
Loss of Top 3b predisposes to spontaneous lymphomagenesis. *(A)* Elevated incidence of lymphomas in Top3b-KO mice. Data are mean ± SEM. *p < 0.05 calculated using two-tailed Student’s t-test. *(B)* Representative splenic structural alterations at 15 months of age. Paraffin-embedded spleen sections stained with hematoxylin and eosin (H&E) and B220, CD3 and F4/80 antibodies show diffuse white pulp expansion in Top3b-KO mice. Scale bar, 1 mm. *(C)* Representative H&E staining of spleens from 15-month-old WT and Top3b-KO female mice. Note dark staining of lymphocytes within the follicular compartment with distinct borders in WT spleen as compared to the pale immature lymphocytes with high rate of mitosis (white arrow) invading into and expanding follicular compartment in Top3b-KO. Higher magnifications of white box areas are shown on the right, Scale bars, 50 μm. *(D)* Representative low magnification images of lymphomas (upper panel) in the spleen and lymph node from 21-month-old Top3b-KO female mouse. Note that neoplastic cells occupy the entire spleen and lymph node. Five-µm paraffin-embedded sections stained with H&E. Scale bar, 1 mm. Microscopic features (lower panel) of lymphoid neoplasms outlined by white boxes in upper panel (*D*) show diffuse expansion of uniform neoplastic cells with distinct border, round nuclei, and scant cytoplasm morphologically resembling lymphoblasts. H&E staining. Scale bars, 100 µm. *(E)* Immunohistochemical staining (IHC) of splenic lymphoma (inset) with anti-B220 and anti-CD3 antibodies. Most neoplastic cells are strongly positive for B220, with a smaller population of CD3^+^ T cells. Scale bar, 100 µm. *(F)* Representative IHC images with lymphocyte-specific (B220, CD3) and proliferation (Ki67) markers in Payer’s patch lymphoma from a 22-month-old Top3b-KO male mouse. Scale bars, 500 µm. Nuclei were co-stained with hematoxylin.

The percentage of mice with tumors greatly increased with age. Of 35 Top3b-KO female mice necropsied at 18-24 months of age, lymphomas were observed in 15 animals versus 3/21 in WT mice (43 % vs. 14.3%, respectively, p < 0.05). Male mice developed lymphomas with similar kinetics, but reduced frequency (Fig. 1*A*) as previously reported for aged C57BL/6 mice (41). There were no differences in histology, anatomical location, and type of tumors between the Top3b-KO female and male mice. Tumors were commonly found in lymph nodes, splenic white pulp, and Payer’s patches of the small intestine (Fig. 1 *D, E* and *F*). In all cases, neoplastic cells displayed lymphoblast-like morphology characterized by large hyper-basophilic nuclei and scant basophilic cytoplasm (Fig. 1 *D*). Immunohistochemical diagnosis confirmed that most lymphoma were composed of the admixed populations of CD3^+^ T and B220^+^ B cells (Fig. 1 *E* and *F*).

Together, these data show that loss of Top3b is associated with enhanced lymphoma development suggesting a tumor suppressor role for Top3b.

### Splenomegaly and inflammatory phenotype in Top3b-KO mice

Systematic investigation of Top3b-KO and WT littermates over time showed an elevation in spleen-to-body weights (SBW) ratios from 6-month of age onwards (Fig. 2*A* and S2). By 18-24 months of age, Top3b-KO female SBWs were on average more than 4-fold higher (p < 0.0001) than in the age-matched WT female littermates (Fig. 2*A*) and 5-fold (p < 0.01) higher than in Top3b-KO male mice (Fig. 2*A* and S2 E). Given a greater and more universal spleen enlargement in the Top3b-KO female mice than in Top3b-KO males, we used females in most of the following experiments.

**Fig. 2.**
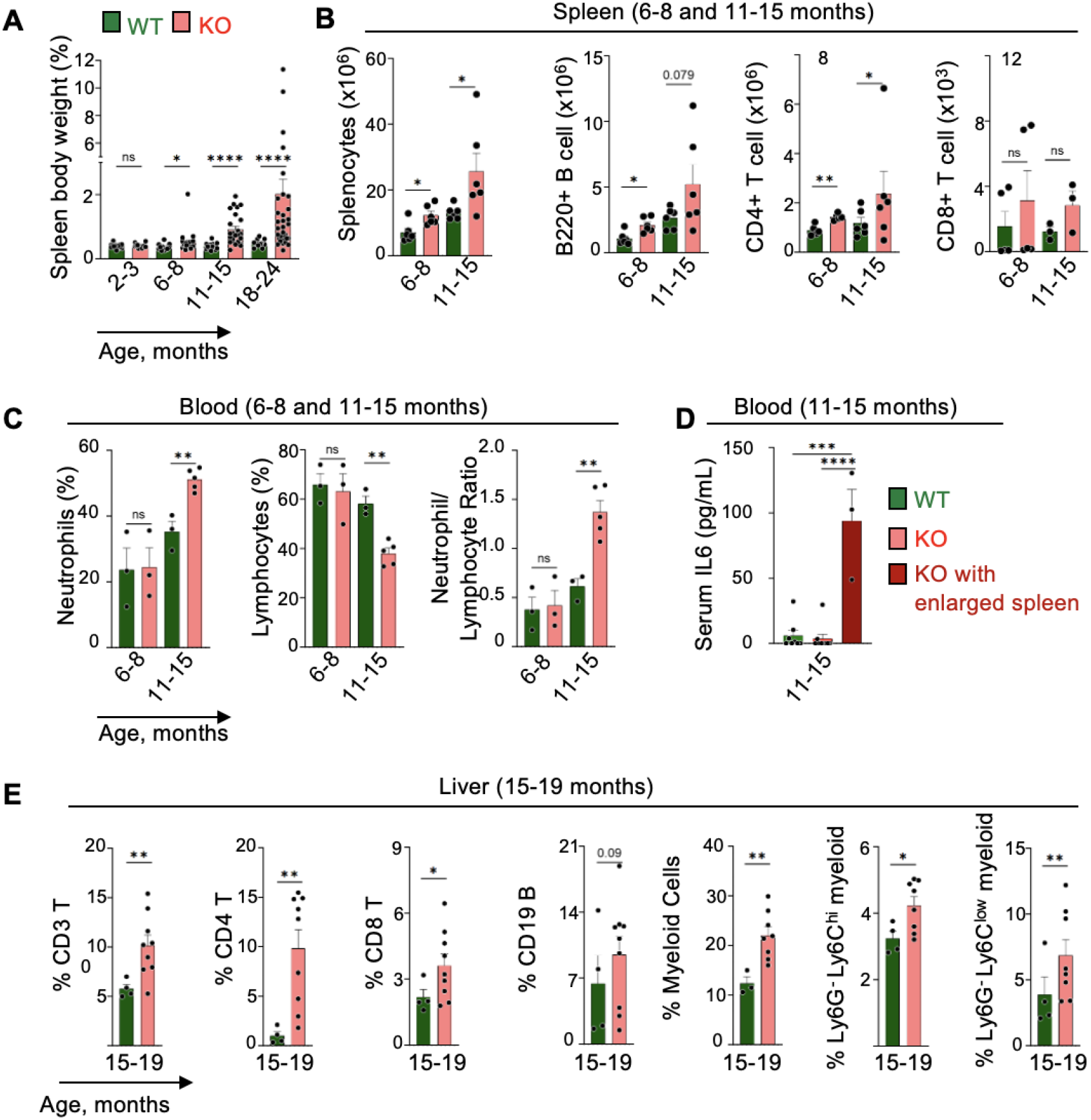
Splenomegaly and immune dysregulation in Top3b-KO mice. (*A*) Spleen-to-body weight ratio in female and male Top3b-KO mice. Mean ± SEM, left to right: n = 15/5/22/4/25/8/33/17 mice. (*B*) Increased splenocytes and Flow cytometry analysis (FACS) of splenic B220^+^, CD4^+^ T and CD8^+^ T cells; Left to right: n = 6/6/5-6/3-5 mice. Data are mean ± SEM. (*C*) Blood analysis showing increase in neutrophils, decrease in lymphocytes and elevation of neutrophil-to-leucocyte ratio in Top3b-KO female mice. Data are mean ± SEM, n =3-5 mice per group. (*D*) Serum interleukin-6 (IL6) levels in 11-15 months female mice: WT (n = 8), Top3b-KO (n = 9). (*E*) Frequency of immune subsets in liver tissues of WT (n=4) and Top3b-KO (n=8) mice at 15-19 months of age. Data are mean ± SEM. p values were determined using unpaired two tailed Welch’s t test. *p < 0.05, **p < 0.01, ***p < 0.001, ****p < 0.0001, ns: non-significant.

In the Top3b-KO mice progressing towards splenomegaly, the total number of splenocytes was increased, reaching more than a 2-fold difference by 11-15 months of age as compared to WT controls (Fig. 2*B*). The higher splenic cellularity was caused by an expansion of B-and T-lymphocytes (Fig. 2*B*) as assessed by flow cytometry analysis (Fig. S3).

Because splenomegaly in rodents can arise from systemic inflammation (42), we performed clinical hematology analyses at different ages. Complete blood cell count analyses revealed a progressive accumulation of neutrophils and concurrent reduction in lymphocytes, resulting in a statistically significant increase in the neutrophil-to-lymphocyte ratio (NLR) (Fig. 2*C*). Top3b-KO females with enlarged spleen also showed a significant elevation in serum interleukin-6 (IL6) level (Fig. 2*D)*. Inflammatory phenotypes observed in spleen prompted us to examine inflammation-associated immune cell infiltration in livers. FACS analysis of livers in Top3b-KO female mice at 15-19 months of age (Fig. S4 A and B) showed a marked accumulation of lymphoid cells as shown by increased CD3^+^ T cells, CD4^+^ T cells, CD8^+^ T and CD19^+^ B cells, and myeloid cells (Fig. 2*E*), implicating a wide range of systemic inflammation (43).

Collectively, these results demonstrate that Top3b loss drives splenomegaly and an inflammatory phenotype.

### Top3b deletion results in genome-wide R-loop accumulation

To gain insight into the molecular mechanisms promoting the tumor predisposition phenotype of Top3b-KO mice, we explored R-loops and the transcriptome of 3-month-old young adult splenocytes (Fig. 3*A*). Given that one of the functions of Top3b is to stabilize the genome by limiting R-loops (8, 14), we compared the abundance of R-loops in WT and Top3b-KO splenocytes by two different methods, R-loop slot-blot (8, 11) and R-Loop Cleavage Under Targets and Tagmentation (R-Loop CUT&Tag) (Fig. 3*A*). A 2-fold elevation of R-loops was found in KO cells compared to WT (Fig. 3 *B* and *C*). The specificity of the R-loop signals observed with S9.6 antibody was confirmed by a treatment of the genomic DNA samples with RNase H before slot-blot analysis (Fig. 3*B*).

**Fig. 3.**
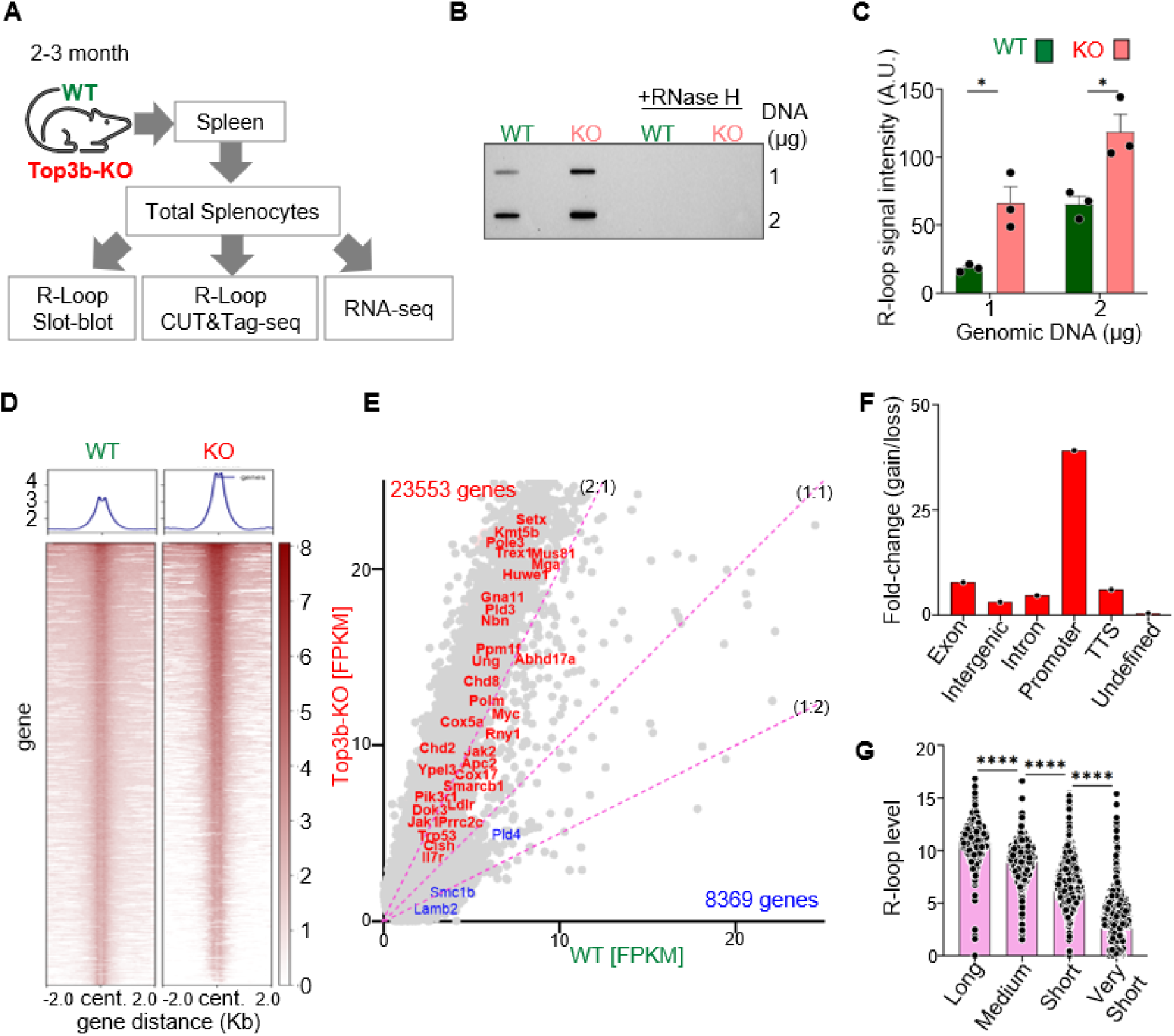
Loss of Top3b results in genome-wide accumulation of R-loops. *(A)* Experimental design. (*B*) Representative slot blot showing increased accumulation of R-loops in Top3b-KO *mouse* splenocytes compared to WT. Genomic DNA was probed with S9.6 antibody. *(C)* Quantitation of 3 independent slot-blot experiments. Data are plotted as means, n = 3 per genotype. Statistical significance was calculated using two-tailed Student’s t-test, *p < 0.05. *(D)* R-loop CUT&Tag-seq signals for WT and KO splenocytes in regions ± 2 Kb from the R-loop centers. Heatmap color intensity represents R-loop CUT&Tag-seq signals level. *(E)* Coverage-plot of normalized R-loop peak signals in 31,922 genes showing R-loops enrichment in 23,553 genes (upper left, read) and depletion in 8,369 genes (lower right, blue) in Top3b-KO mice. FPKM, Fragments Per Kilobase Million. (*F*) Distribution of R-loop peak intensity expressed across different genomic features. (*G*) Average R-loop signal intensity is positively correlated with gene length. 31,922 genes were divided into four different quartiles: Very short (< 0.1 kb), Short (≥ 0.1 kb and < 1 kb), Medium (≥ 1 kb and < 10 kb) and Long (≥ 10 kb), and distribution of average R-loop signal intensity in each quartile is plotted. p values were determined using the Mann Whitney test. ****p < 0.0001.

Genome-wide native R-loop profiling by R-Loop CUT&Tag confirmed that Top3b-KO splenocytes displayed an increase in R-loop peak intensities across 72 % of the total R-loop peaks (Fig. S5). The average peak intensity was elevated approximately 2-fold in Top3b-KO splenocytes as shown in the metaplot of Figure 3*D* (Upper panel) and the heatmap of Figure 3*D* (lower panel). The raw R-loop peak intensities were then assigned to the corresponding genes. In total, we analyzed R-loops across 31,922 genes and found increased R-loop accumulation across 73.8 % of all gene loci in Top3b-KO splenocytes (Fig. 3*E*). These results agree with the R-loop slot-blot data (Fig. 3 *C* and *E*). Differential peak analysis (FC > 1.2 or FC < -1.2, p-value < 0.05), showed the highest R-loop signal enrichment at the promoter regions (Fig. 3*F*).

To address the relationship between gene length and R-loop levels, the 31,922 genes were divided into four different quartiles: Very Short, Short, Medium and Long. The average R-loop signal intensity in each quartile was directly related to the gene length (Fig. 3*G*) indicating that longer genes accumulate more R-loops in the Top3b-KO splenocytes.

Collectively, these results demonstrate that Top3b prevents genome wide accumulation of R loops with the greatest impact at the promoter regions and in long genes.

### Top3b deletion is associated with dysregulated immunotranscriptome

RNA-seq analysis of Top3b-KO splenocytes performed in parallel with the R-loop experiments identified 1,371 (Dataset 01) differentially expressed genes (DEGs) using the Linear Model for Microarray Data (limma) (44). Among these, 509 (32%) were upregulated and 862 genes (63%) were downregulated (Fig. 4 *A* and *B*). Functional annotation of the 1,371 DEGs using Ingenuity Pathway Analysis (IPA) revealed a significant enrichment of several immune signaling pathways including “Interleukin-7 signaling”, “IL-6 Signaling” and suppression of “Dendritic Cell Maturation”, “Neutrophil degranulation (Fig. 4*C* and Dataset 01). These data support that Top3b-loss is associated with immune dysregulation.

**Fig. 4.**
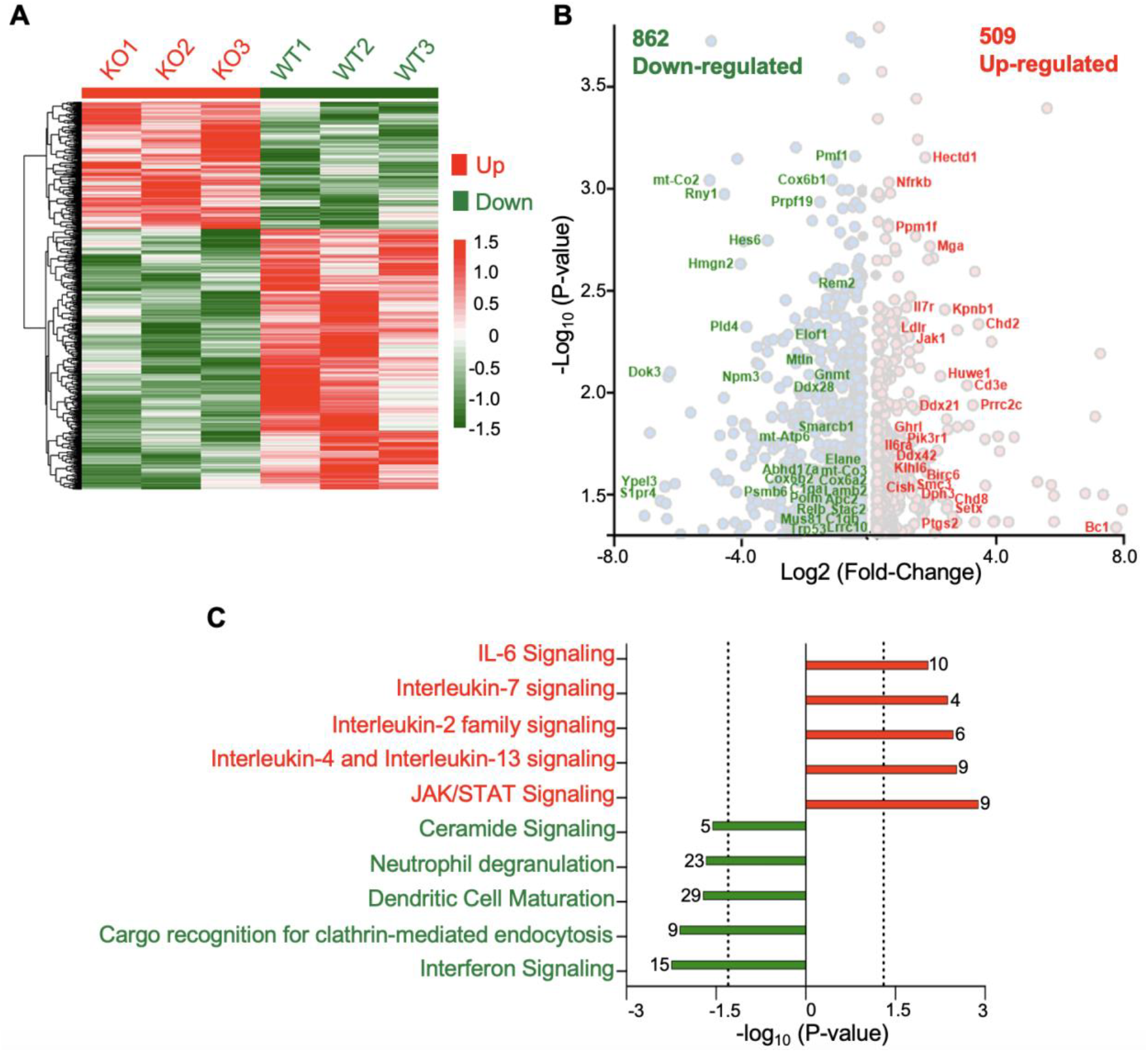
Top3b loss leads to dysregulation of the immunotransciptome. (*A*) Heatmap showing distribution of RNA signal across differentially down-regulated (862) and up-regulated (509) genes in three independent RNA-seq experiments in Top3b-KO splenocytes. (*B*) Volcano plots showing some of the differentially expressed genes (DEGs) in Top3b-KO splenocytes. DEGs were analyzed using LIMMA. *(C)* Functional annotation of the significantly altered immune pathways of DEGs (1,371) determined by Ingenuity Pathway Analysis.

To evaluate the relationship between R-loop elevation and transcriptomic alterations in Top3b-KO splenocytes, R-loops levels were plotted against RNA-expression (Fig. 5*A*). Lack of correlation indicates that R-loop elevation in Top3b-KO splenocytes does not impact RNA-expression (Fig 5*A*). The lack of correlation was further demonstrated by Integrative Genome Viewer (IGV) tracings of three representative DEGs including *Myc, Elof1*, and *Ppm1f* (Fig 5B). The graphs showed that the R-loop signals were enhanced in the absence of Top3b, while the transcript levels were relatively similar. Of note, when we plotted the average R-loop levels among the DEGs, we observed that highly expressed genes accumulate more R-loops than the downregulated genes (p < 0.001) (Fig. S6A). Again, the abundance of R-loops in the DEGs was dependent on the gene length (Fig. S6 B and C).

**Fig. 5.**
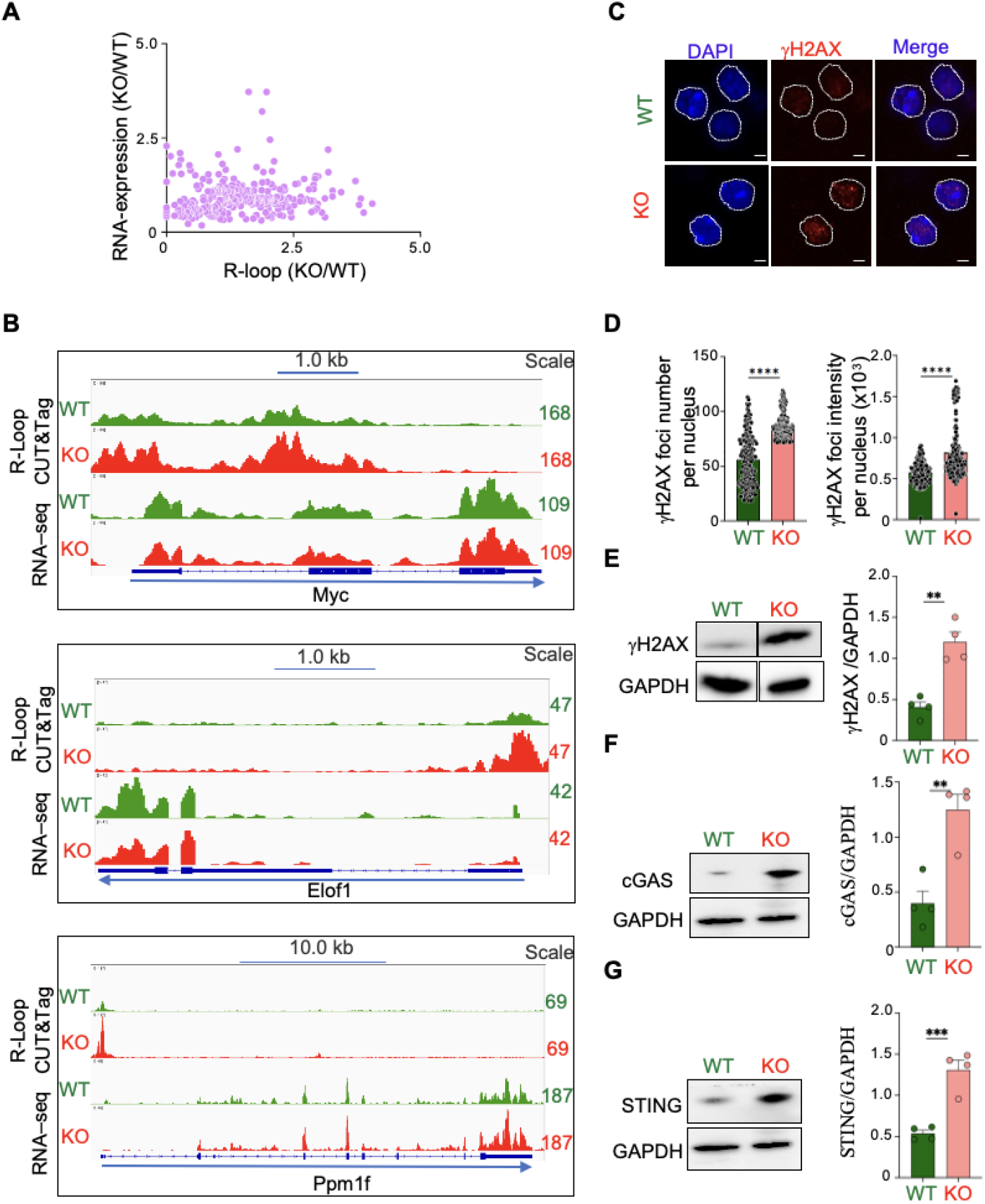
Increased DNA breaks and cGas-Sting protein levels in Top3b-KO splenocytes. *(A*) Lack of correlation between R-loop levels and RNA-expression. *(B)* Examples of R-loop signals (IVG tracing) in representative genomic regions determined by CUT&Tag-seq in WT (green) and KO (red) splenocytes. R-loop signals were group auto-scaled with the Top3b; scales are indicated at right. (*C*) Representative image of immunofluorescence staining of 3-month-old female mouse splenocytes with anti-gamma-H2AX (γH2AX) antibody. *n* = 3 mice per genotype. Nuclei were co-stained with DAPI and outlined with white lines. Scale bar, 10 μm. *(D)* Quantification of γH2AX foci number and foci intensity shown in (*C*). 200 nuclei were analyzed by Random-Forest ML pixel-classifier per each mouse. Data are means ± SEM; ****p < 0.0001, unpaired, two-tailed Student’s t-test. *(E-F)* Detection and quantification of γH2AX (E), cGas (F) and Sting (G) by Western blotting. GAPDH was used as loading control. n = 4 mice per genotype. Data are means ± SEM; **p < 0.01, ***p < 0.001 unpaired, two-tailed Student’s t-test.

These data demonstrate that R-loops do not directly affect genome-wide transcripts apart from a few highly expressed DEGs and long genes.

### DNA breaks and innate immune activation in Top3b-deleted splenocytes

To assess whether R-loop accumulation is associated with increased DNA damage (45), we performed immunofluorescence staining of 3-month-old young adult splenocytes with γH2AX antibody, a well-established sensitive molecular marker of DSBs (46). Figure 5*C* shows increased γH2AX in Top3b-KO splenocytes. Quantitation by Random-Forest ML pixel-classifier showed a significant increase in γH2AX foci number (Fig. 5*D*) as well as signal intensity (Fig. 5*D*) in Top3b-KO as compared to WT splenocytes controls. Western blotting experiments confirmed elevation of the γH2AX signal in Top3b-KO splenocytes (Fig. 5*E*).

Since aberrant R-loop formation is known to cause cytoplasmic accumulation of DNA/RNA/RNA-DNA hybrid species leading to activation of the cGAS-STING signalling pathway (47, 48), one of the major cytosolic DNA sensors in mammalian cells with diverse functions in host immunity and production of pro-inflammatory cytokines (49, 50), we performed Western blotting of cGAS (Fig. 5*F*) and Sting (Fig. 5*G*). Both cGAS and Sting proteins levels were significantly increased in KO splenocytes as compared to WT.

These results suggest a previously unappreciated role of Top3b in preventing genomic breaks and innate immune activation.

### Top3b-deficiency promotes syngeneic EL4 lymphoma growth

Because elevated activation of the cGAS-STING pathway has been associated with peripheral T-cell lymphomas (51) and we found the Top3b-KO mice to be immune-compromized, we used a lymphoma syngeneic model (Fig. 6*A*) and compared tumor development in Top3b-KO and WT mice. Since EL4 cells were derived from a malignant T-cell lymphoma induced in a C57BL/6 inbred mouse strain, this model allowed us to compare the impact of a functional (WT) and Top3b-deficient microenvironment on lymphoma growth.

**Fig. 6.**
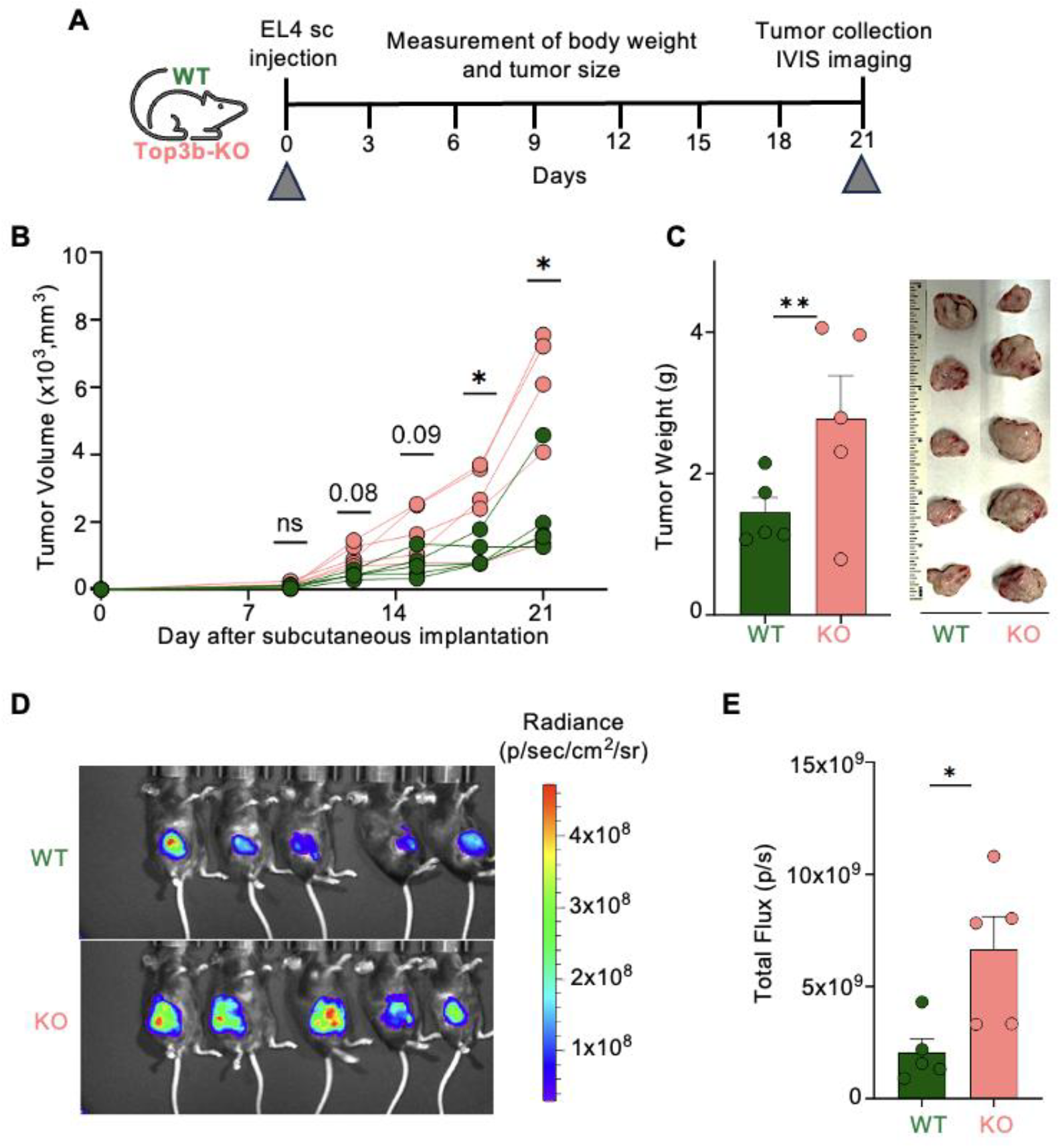
Top3b-deficiency promotes syngeneic growth of EL4 lymphoma. *(A)* Experimental scheme. 10,000 EL4 tumor cells stably transfected with a luciferase reporter gene were injected subcutaneously (s.c.) into the right flank of 3-month-old female mice (*n* = 5 for each genotype). Data are representative of two biological independent experiments with similar results. *(B)* Kinetics of tumor growth. Tumor size was determined using a digital caliper twice a week. Data were analyzed by unpaired, two-tailed Student’s t-test. *p < 0.05, ns, not significant. *(C)* Tumor weights 21 days after EL4 transplantation. Photographs on the right show differences in tumor size at the end of experiments. Data are means ± SEM; **p < 0.01, two-tailed Student’s t-test, ns, not significant. *(D)* Representative bioluminescence images 21 days after transplantation of EL4 cells. *(E)* Quantification of bioluminescence imaging as total flux of photons per second. Data are means ± SEM; *p < 0.05, two-tailed Student’s t-test, ns, not significant.

WT and Top3b-KO mice were challenged with subcutaneous injections of 10,000 EL4 cells stably expressing a luciferase reporter gene. EL4 lymphoma cells were readily engrafted both in WT and Top3b-KO recipients. Notably, in Top3b-KO mice, tumors grew faster and formed larger tumors than in WT mice (Fig. 6*B*). As a result, syngeneic EL4 lymphomas in Top3b-KO recipients reached an experimental tumor size limit of ≤ 20 mm at 21 days after transplantation, significantly earlier than in the WT control mice (Fig. 6*C*). Bioluminescence imaging on day 21 showed on average a 3-fold increase in tumor mass in Top3b-KO recipients (Fig. 6 *D* and *E*). The transplantation caused no toxicity as evaluated by the mouse weights (Fig. S7*A*).

Collectively, these data demonstrate that loss of Top3b promotes lymphoma growth.

## Discussion

Herein, we report that genetic inactivation of Top3b promotes syngeneic lymphoma growth in young adult mice and accelerates spontaneous lymphomagenesis in aging mice (Figs. 1 & 6). At the molecular level, we provide evidence that loss of Top3b leads to aberrant accumulation of R-loops (Fig. 3), genomic DSBs (Fig. 5) and immune dysregulation (Fig. 2, 4, 5). We propose that genetic instability in combination with impaired immune function renders Top3b-KO mice susceptible to lymphomagenesis in addition to the know functons of Top3b in ensuing normal neurologenesis and synaptic plasticity (10, 52, 53).

Lymphomas are among the most common naturally occurring malignancies in many strains of mice with a higher incidence in females (41, 54-56), and a major cause of mortality in C57BL/6 mice (41). In agreement with published studies (36), we find that Top3b-KO mice display a shorterned life span, which we attribute, at least in part to increased lymphoma burden (Fig. S1A), especially in females mice (Fig. 1).

Upon aging, we find that Top3b mice exhibit features of immune system dysfunctions including splenomegaly, inflammatory cell infiltration in different organs, and a higher frequency of ulcerative dermatitis (Fig. S1 *D* and *E*), which is consistent with the prior report from Shaw and Wang (40). At the molecular levels, we find that Top3b deficiency causes a statistically significant increase in the neutrophil-to-lymphocyte ratio (NLR) (Fig.1 *E*), a marker of systemic inflammation and of diseases including infectious and autoimmune disorders (57), schizophrenia (58) and cancer (59, 60). These findings highlight a critical role of Top3b in preventing systemic inflammation and lymphomagenesis.

Our study further establishes that Top3b acts as a main suppressor deleterious R-loops driving genomic instability *in vivo* (3). Top3b loss-associated accumulation of R-loops and subsequent DNA damage were reported in human renal cancer (14). This finding is consistent with the observed elevation of R-loops and DSBs at the c-Myc and Ig heavy chain (Igh) loci in mice lacking Tdrd3, the scaffolding partner of Top3b (2, 15). Impaired Top3b activity has also been related to mammary gland cancer (61), bilateral renal cancer (14, 62), lymphoma (63) and melanoma (64). Characterization of Top3b-KO splenocytes isolated from 2-3-month-old mice revealed accumulation of R-loops with a parallel increase in DNA damage reflected by the presence of γH2AX Analysis of R-loops across the entire genome demonstrates enrichment of R-loop signals at promoter regions (Fig. 3*F*).

Among the hallmark features of impaired processing of R-loops is activation of cGas/Sting-mediated inflammatory response through exposure to self-DNA/RNA/DNA-RNA hybrid (45, 47). Accordingly, we found increased cGas/Sting protein levels Top3b-KO splenocytes (Fig. 5 *F* and *G*). Furthermore, pathway analysis of the DEGs showed a statistically significant enrichment of the major inflammatory pathways, including but not limited to Th1 Pathway, interleukin-3, interleukin-5 and GM-CSF signaling, Acute Phase Response (APR) signaling and interleukin-6 signaling (Fig. 4 *C* and Dataset S01).

In summary, our findings demonstrate that Top3b-KO mice develop a complex phenotype linking Top3b deficiency with R-loop mediated genomic instability that triggers immune dysregulation and lymphoma-prone phenotype of Top3b-KO mice.

## Materials and Methods

### EL4 LUC xenograft model in WT and Top3b-KO C57BL/6mice

2–3-month-old females WT and Top3b-KO mice were used in this experiment. All procedures were performed in accordance with the guidelines of the Animal Care and Use Committee (ACUC) of the National Cancer Institute, NIH. The cells were negative for mycoplasma and all the viruses tested. EL4 cells (1×10^4^ cells/mouse) were subcutaneously transplanted into the right flank of experimental mice in a 1:1 mix of HBSS and Matrigel. Tumor size and mouse weight were assessed every three days until day-21. Tumor volume was calculated by the formula: volume = 0.5 × length × (width)^2^. At day-21 mice were injected with D-luciferin (150 mg/kg body weight) at the right quadrant of the abdomen intraperitoneally for bioluminescence imaging. Images were taken 10 min after injection using a Xenogen IVIS System. Signal intensity quantification was performed using the Living Image software (Xenogen).

### Data, Materials, and Software Availability

All study data are included in the article and/or supporting information. All original sequencing data will be submitted in GEO before final publication.

## Acknowledgements

We thank CCR Genomics Core, NCI-Bethesda, Maryland for helping with CUT&Tag-seq and RNA seq (splenocytes) library preparation and sequencing, and Dr. Baktiar Karim, MHL, NCI, Frederick and Tyler Peat, D.V.M., Ph.D., NCI for the histopathological evaluation of slides. We thank Building 37, NCI, NIH animal facility staffs Devorah Gallardo, Kristin Killoran and Alilin Miafor their unwavering supports for this study. We are grateful to Andy Tran (Confocal Microscopy Core Facility, CCR, NCI, NIH) for support with confocal microscopy analyses and Dr. Jennifer Dwyer for scanning IHC slides. Our studies are supported by the Center for Cancer Research, the Intramural Program of the National Cancer Institute, NIH, Bethesda, MD 20892 (Z01 BC 006161-17 and Z01 BC 006150-19).

