## Supporting Information for "Topoisomerase IIIβ protects from tumorigenesis and immune dysregulation"

**Affiliations**

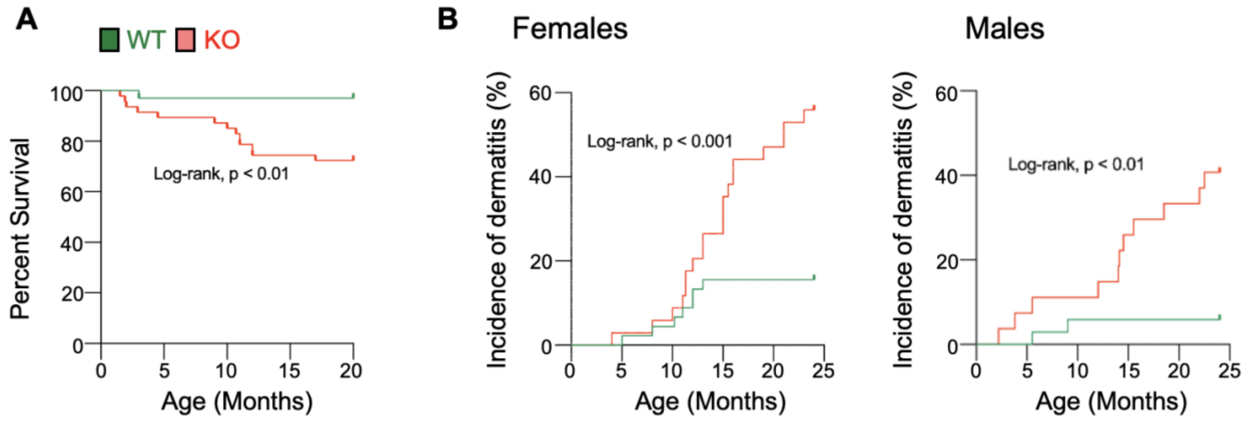


**Fig. S1. Reduced lifespan and increased frequency of ulcerative dermatitis in Top3b-KO mice.** (Related to Fig. 1). (A) Kaplan-Meyer survival curves. n = 35 (WT) and n=48 (Top3b-KO). (B) Incidence of ulcerative dermatitis in female (middle panel) and male (right panel) mice. Number of mice in females: n = 45 (WT) and 34 (Top3b-KO) and males: n = 34 (WT) and 27 (Top3b-KO).

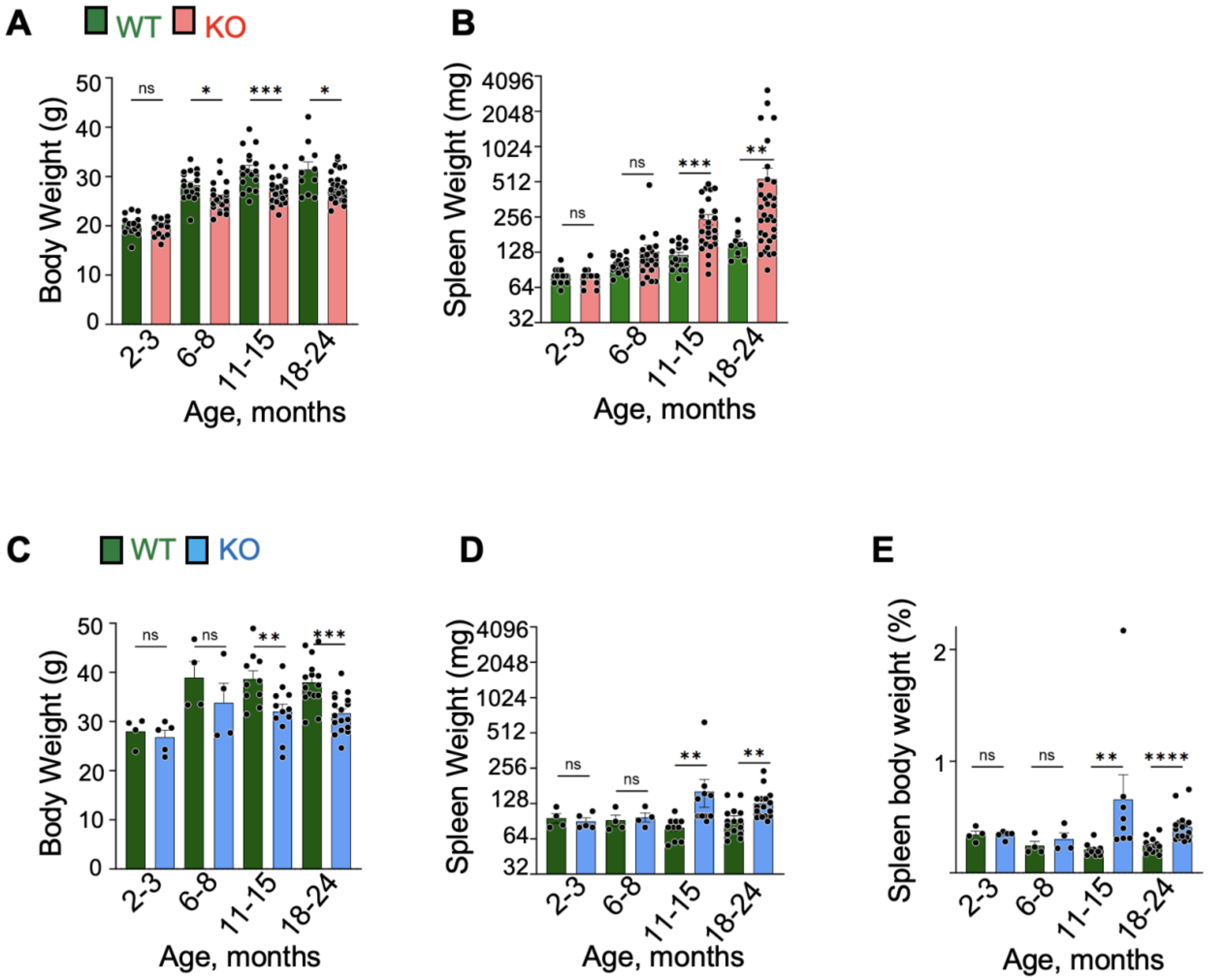


**Fig. S2. Increased spleen weight and reduced body weight in KO mice.** (Related to Fig. 2). (A) Body weight of female mice. Mean ± SEM, left to right: n=15/15/19/22/17/25/11/33 mice. (B) Spleen weight in female mice. Mean ± SEM, left to right: n = 15/15/19/22/17/25/11/33 mice. (C) Body weight of male mice. Mean ± SEM, left to right: n = 4/5/7/11/23/28/25/42 mice. (D) Spleen weight in male mice. Mean ± SEM, left to right: n = 4/5/4/4/10/12/15/17 mice. (E) Age-dependent increase in spleen-to-body weight percentage in male mice. Mean ± SEM, left to right: n = 4/5/4/4/10/8/15/17 mice. p values were determined using Welch's t test. *$p < 0.05$, **$p < 0.01$, ***$p < 0.001$, ****$p < 0.0001$, ns: non-significant.

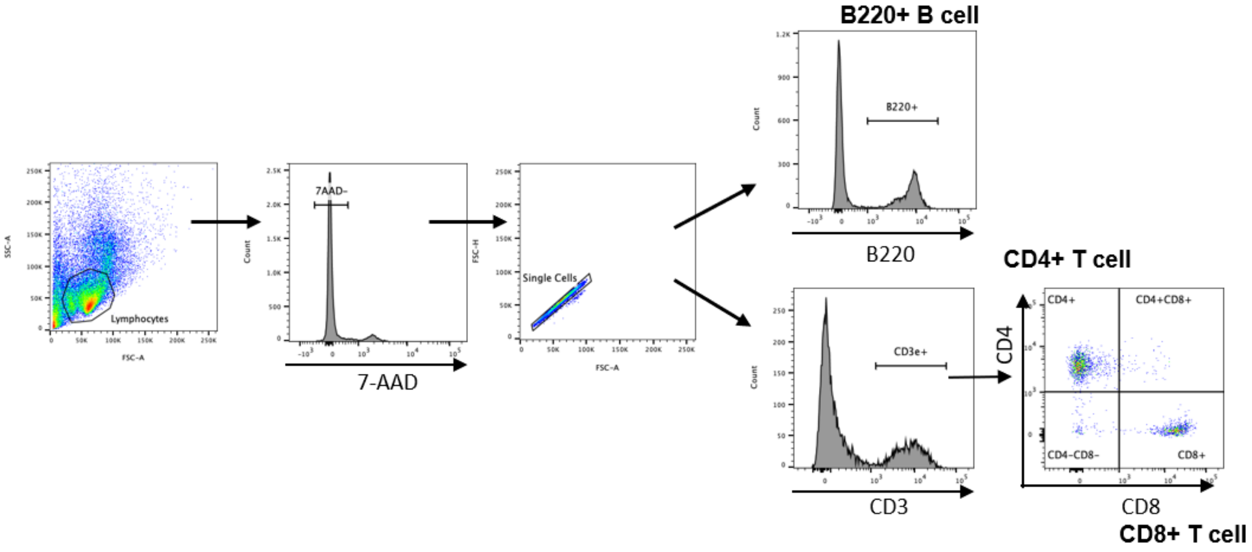


**Fig. S3. Gating strategy for FACS analysis of splenic lymphocytes isolated from WT and Top3b female mice at 6-8 and 11-15 months of age.** (Related to Fig. 2). Splenocytes were isolated from 3 mice per genotype and analyzed individually.

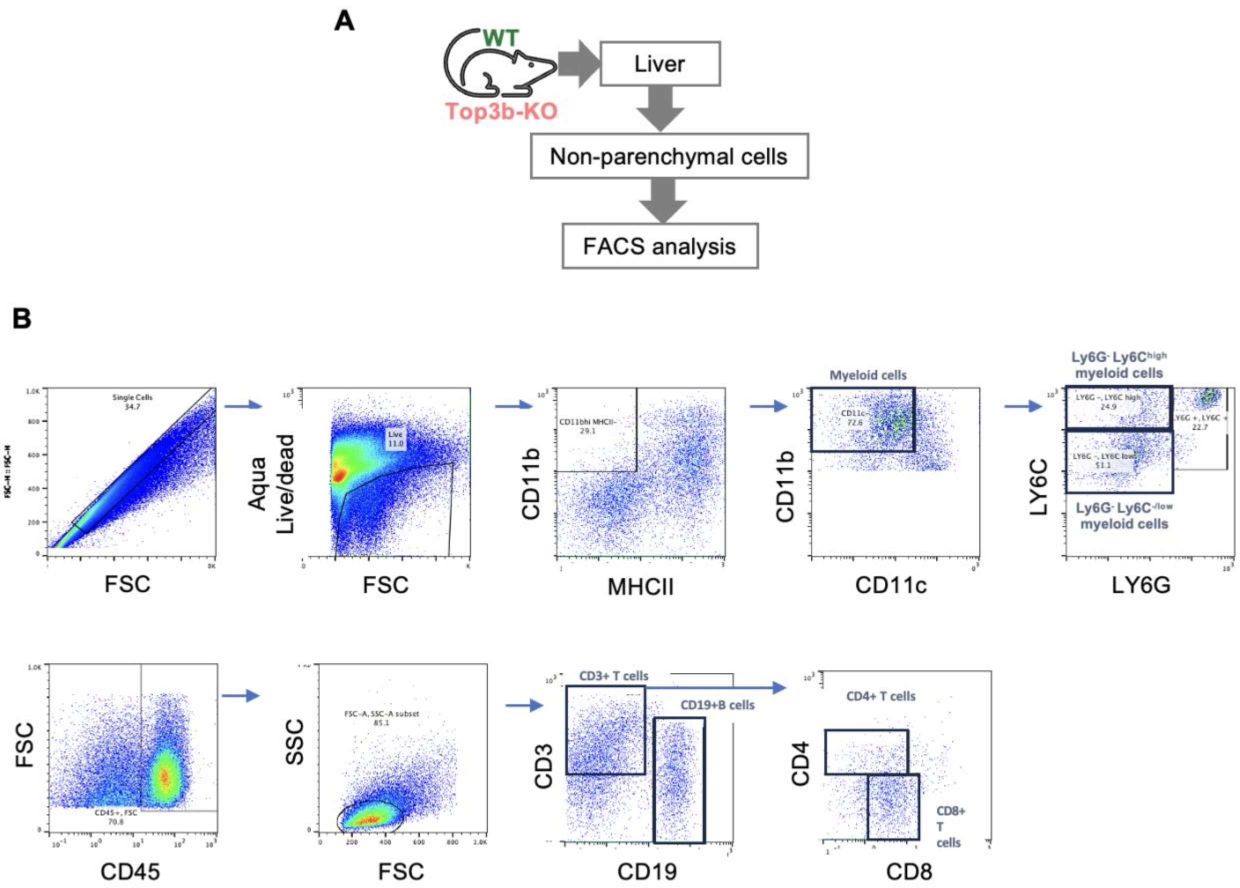


**Fig. S4. Top3b deficiency increases immune cell infiltration into liver.** (Related to Fig. 2). (A) Schematic diagram for analysis of immune cells infiltrating livers of 15-19-month-old female mice. (B) Gating strategy for FACS analysis of immune cells. Single cells dissociated from liver tissues were first gated for CD11b+ MHC class II-. CD11b positive and CD11c- cells were then gated for Ly6G- and Ly6C high or low. CD45+ cells were first gated for CD3+ cells, CD3+ cells were further sub-gated for CD4+ and CD8+ T cells.

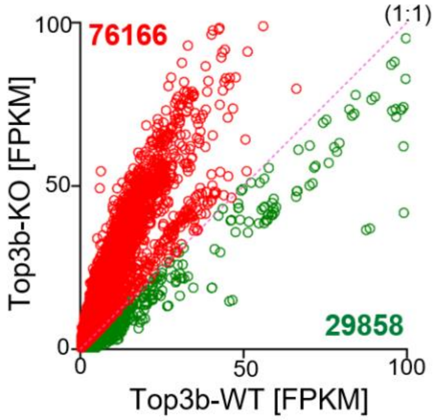


**Fig. S5. Top3b-loss enhances genome-wide R-loop peaks in Top3b-KO splenocytes.** (Related to Fig. 3)**.** Coverage-plot Fragments Per Kilobase of transcript per Million mapped reads of normalized R-loop peaks' signal. Total number of peaks plotted, n=106024. Red peaks (n = 76166) were enriched, and green peaks (n= 29858) were depleted in Top3b-KO cells, respectively.

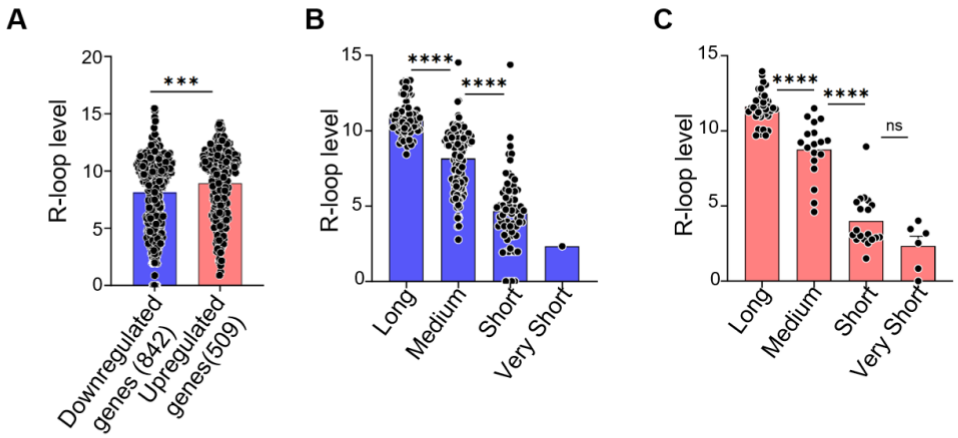


**Fig. S6. RNA expression and R-loop levels.** (Related to Fig. 4).
(A) Average R-loop signal intensity of differentially expressed down-regulated and up-regulated genes in Top3b-KO splenocytes compared to WT controls. (B) Average R-loop signal intensity of 862 genes that were downregulated in Top3b-KO compared to WT splenocytes is positively correlated with gene length. Genes were divided into four different quartiles: Very short (<0.1 kb), Short (≥ 0.1 kb and < 1 kb), Medium (≥ 1 kb and < 10 kb) and Long (≥ 10 kb), and distribution of average R-loop signal intensity for all the genes in each quartile was plotted. (C) Average R-loop signal intensity of 509 genes that were upregulated in Top3b-KO compared to WT splenocytes is positively correlates with gene length. Genes were divided into four different quartiles: Very short (<0.1 kb), Short (≥ 0.1 kb and < 1 kb), Medium (≥ 1 kb and < 10 kb) and Long (≥ 10 kb), and distribution of average R-loop signal intensity for all the genes in each quartile are plotted. p values were determined using Mann Whitney test (A-C). *$p < 0.05$, ****$p < 0.0001$ unpaired, ns, non-significant.

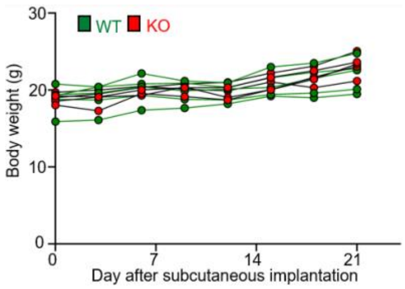


**Fig. S7. Changes in body weight after EL4 implantation.** (Related to Fig. 6). There were no differences in body weight between WT and Top3b-KO female mice after subcutaneous implantation of EL4 lymphoma cells.

### Supplementary materials and methods

#### Cell line

EL4 cell line purchased from ATCC (cat # TIB-39) and was grown in Dulbecco's modified Eagle's medium (Life Technologies, Carlsbad, CA) supplemented with 10% Fetal Bovine Serum (Gemini, West Sacramento, CA, 100-106), 1% penicillin-streptomycin (ThermoFisher Scientific, 15140122) and 55 mM 2-mercaptoethanol at 37°C in humidified 5% CO2 chamber. All experiments were performed within 25 passages from thawing, and cell lines were routinely tested for mycoplasma (Lonza, LT07-318) contamination.

#### Mouse model

The Top3b-KO mouse line (36) (a gift from Dr. Albert C Shaw, Yale University) was rederived before introducing into our facility. The mouse colony was maintained in an outbred C57BL/6 background under standard housing conditions (21 ± 1°C) with artificial light (12:12-hr light: dark cycle). Mice were kept in isolated cages. To maintain a specific pathogen-free environment in the cage, controlled ventilation was performed using HEPA-filters. Mating of individual pairs was initiated when the animals reached 8 weeks of age, and the litter size from each individual pair was monitored over a period of 8–10 months. All mice were genotyped at weaning by commercial vendor (Transnetyx, Cordova, TN).

#### Hematoxylin and eosin (H&E) and Immunocytochemistry staining

All animal work was performed in compliance with relevant regulatory standards and was approved by the Animal Research Committee of National Cancer Institute, NIH Animal Care and Use Committee. To prepare paraffin block for histological analysis, organs were fixed overnight in 10 % neutral buffered formalin, embedded in paraffin and cut into 5-μm slices by microtome for either H&E staining for routine histology or immunostaining staining with antibodies against B-cell, T-cell, and macrophages with All immunostainings were performed in house or by Histoserv, Inc. (Gaithersburg) and/or Molecular Histopathology lab, NCI (Frederick) using standard procedure.

#### Genetic modification of the EL4 cell line expresses the Luciferase gene

Lentivirus was produced by transfection of HEK-293T cells with lentiviral plasmid of interest, Pax2 (Adgene 35002), and VSVG (Adgene 14888). Lentiviral plasmid pLenti CNV V5-LUC Blast (Adgene 21474) for luciferase. The viral production was performed according to the NIH ACUC guidance. The HEK-293T cells were seeded in DMSE+10% FBS on day-1trasnfected using turbofectin 8.0 on day 0 (OrgiGene TF81001) and the virus was collected after 48-72 hours of post transfection. The luciferase viral transduction was achieved by seeding 1-2 million EL4 cells with one ml of viral soup, along with 10 µl/ml of Polybrene (Thermo Scientific), by centrifuge at 1300g for 2 hours at $24^0$C and followed by incubation overnight at $37^0$C in a $CO_2$ incubator. Cells were washed and cultured in EL4 cell media supplemented with 5µl/ml blasticidin for a week. Luciferase cells were reduced to maintenance of 1µg/ml and eventually taken off blasticidin entirely.

**R-loop Dot-Blot**

For R-loop detection by slot-blot, genomic DNA was extracted from isolated splenocytes using protocol as described previously (77-79). Briefly, cells were lysed in TE buffer containing SDS and proteinase K (at 37°C overnight, Invitrogen™ Proteinase K Solution (20 mg/mL), phase separated using phenol/chloroform/isoamyl alcohol (25:24:1), ethanol precipitated and resuspended in TE buffer. Genomic DNA was digested using cocktail of restriction enzymes (HindIII, SspI, EcoRI, BsrGI and XbaI; 30 U each), treated with RNase T1 (2 units; ThermoFisher Scientific Cat# EN0542) and shortcut RNase III (2 units; New England Biolabs; Cat# M0245L) and again purified by phenol/chloroform/isoamyl alcohol (25:24:1) extraction. Increasing concentrations of genomic DNA were spotted on a nitrocellulose membrane, crosslinked with UV light (120 mJ/cm2)), blocked with PBS-Tween (0.1%) buffer and 5% non-fat milk (Room temperature for 1hr) and incubated with mouse S9.6 antibody (1:500 dilution, overnight at 4°C, Millipore Sigma, Cat# MABE1095). After washing with PBS-Tween (0.1%), membrane was incubated with HRP-conjugated anti-mouse secondary antibody, washed, and developed with ECL techniques. In case of RNase H (New England Biolabs, Cat# M0297L) treated control, 10 µg genomic DNA was pre-incubated with 20 U of RNase H for three hours at 37°C (77).

**Western Blotting**

Splenocytes were lysed in 100 µL sodium dodecyl sulfate (SDS) buffer containing 25 mM Tris–HCl (pH 6.5), 1% SDS, 0.24 mM β-mercaptoethanol, 0.1% bromophenol blue and 5% glycerol and cOmpleteTM, Mini, EDTA-free protease inhibitor cocktail (Roche, Cat# 11836170001). Whole-cell extracts were separated by SDS-PAGE gel, transferred onto polyvinylidene difluoride (PVDF) membranes, and blocked in 5% skimmed milk dissolved in 0.1% Tween-20 in phosphate buffer saline (PBS). Membranes were incubated with primary antibodies overnight at 4 °C followed by washing with 0.1% Tween-20 in PBS. Primary antibodies used in this study are as follows: rabbit polyclonal anti-γH2AX (dilution 1:1000, Cell Signaling, Cat # 9718), rabbit monoclonal anti-STING (dilution 1:1000, Cell Signaling, Cat # 13647), and rabbit mAb anti-GAPDH (dilution 1:2000, Cell Signaling Technology, Cat# 2118, Clone 14C10). Membranes were incubated with anti-rabbit IgG ECL, HRP conjugated (dilution 1:4000, GE Healthcare, Cat# NA9340) at rt for 1 h and washed thrice and signals were detected by ECL chemiluminescence reaction (SuperSignal™ West Femto Maximum Sensitivity Substrate, Thermo Scientific, Waltham, MA, Cat# 34095).

**R-loop CUT&Tag sequencing**

The Cut and Tag protocol was carried out following Zhang et al (80). Briefly, 2 μl 2XCB (0.1 M Tris pH 8.0; 0.3 M NaCl; 0.1% Triton X-100; 25% Glycerol), 2 μl pA-Tn5 tranposome (at 4 μM), and 2 μl S9.6 antibody were mixed in a microcentrifuge tube (mixture A) and kept on ice. One million cells were harvested and transferred to a clean 1.5 mL tube. Cells were washed once with 1 ml of PBS, suspend in 1 mL of 1X CB, and incubated on ice for 10 min. The cells were spun down, suspend in 1 mL of 1X CB. An aliquot of 50 μl was added to mixture A, incubated at room temperature for 60 min with rotation. 500 μl of wash buffer (50 mM Tris pH 8.0; 150 mM NaCl; 0.05% Triton X-100) was added to each tube, centrifuged. The pellets were washed two more time with 500 μl of wash buffer and then suspended in 100 μl of wash buffer. 1 μl of 1 M $MgCl_2$ was added to start the transposome reaction, incubated at 37°C for 60 min. 4 μl of 0.5 M EDTA, 2 μl of 10% SDS and 1 μl of 20 mg/mL Proteinase K were added, and incubated at 55 °C for 60 min. The DNA was purified with ChIP DNA clean & concentrator (Zymo Research). Library was constructed following manufacture's protocol (Illumina) and sequenced on Next-seq.

**Total RNA sequencing**

Total RNA was extracted by TRIzol (Invitrogen), and then purification using PureLink RNA Mini Kit (Cat No. 12183018A; Invitrogen) with DNase treatment (QIAGEN). 2100 Expert (Agilent) was used for RNA quality and quantity assessment. RIN (the RNA Integrity Number) of all samples were above 9.0.

Library was prepared for splenocytes using NEBNext rRNA Depletion Kit v2 (Human/Mouse/Rat) (NEB #E7400) and NEBNext Ultra II Directional RNA Library Prep Kit from Illumina (NEB #E7760). For PCR amplification (13 cycles) NEBNext Multiplex Oligos for Illumina (96 Unique Dual Index Primer Pairs, NEB, E6440) were used. Dual index (101x8x8x101) pair-end sequencing of the final 750 pM library was performed using the Illumina NextSeq2000 P2 Reagents (200cycles) on the Illumina NextSeq2000 instrument. The total number of reads was 550M.

Biotinylated, target-specific oligos combined with Ribo-Zero ribosomal RNA (rRNA) removal beads were used for the removal of rRNA library construction for splenocytes. Illumina TruSeq Stranded Total RNA Library Preparation kit and paired-end sequencing of 6 pooled RNA-seq samples was performed on HiSeq. Following RNA fragmentation and cleavage, small pieces are converted to complementary DNA (cDNA) with reverse transcriptase and random primers and, consequently $2^{nd}$ strand cDNA synthesis with DNA Polymerase I and RNase H. A standard Illumina library prep with end-repair, adapter ligation and PCR amplification being performed to give you a sequencing ready library from the double-strand cDNA as input. After that, purified product was quantified using qPCR followed by cluster generation and pair-end sequencing on HiSeq.

**Data Preprocessing and Bioinformatics Analysis**

R-loop CUT&Tag sequencing fastq files for WT and Top3b-KO were generated from splenocytes of young female mice. Sequencing quality was assessed using FASTQC (version 0.12.1), and duplicate reads were identified using Picard (version 3.2.0). After duplicates were removed, the fastq files were aligned to the mouse reference genome (mm39) using BWA aligner (version 0.7.17). Peak calling was conducted with MACS2 (version 2.2.7.1), and normalized bigwig files were produced using the BAMscale "scale" module (81). Additionally, coverage of aligned BAM files was calculated with BAMscale "cov" (version 0.0.6), applying the FPKM normalization method. Peaks annotations and differential peak analysis was performed using HOMER software (version 5.1) using "annotatePeaks.pl" and "getDifferentialPeaks" modules. Average signal intensity heatmaps were generated using plotHeatmap module and read coverage of WT and Top3b-KO bigWig files calculated using "multiBigwigSummary" module of deepTools (version 3.5.5) (82).

In addition, RNA-seq fastq files from WT and Top3b-KO female mouse splenocytes were aligned to mm39 reference genome using STAR aligner (version 2.7.10b) (83). SAMtools (v 1.17) used to sort, and index aligned BAM files. Read counts per gene were computed with the STAR "–quantMode GeneCounts" option, and raw reads normalized to RPKM (Reads Per Kilobase per Million mapped reads) values. Differential gene expression analysis was conducted using "limma" package [https://bioconductor.org/packages/release/bioc/html/limma.html] in R (version 4.2.3). Differentially expressed genes (DEGs) were used to perform pathway analysis using Ingenuity Pathway Analysis (IPA) software (Ingenuity Systems; Qiagen China Co., Ltd.).

#### γH2AX Immunostaining

For γH2AX detection, total splenocytes (0.5 X $10^6$ cells) were collected using Cytospin and fixed with 4 % PFA in PBS for 15 minutes at room temperature (RT) followed by permeabilization for 15 minutes with Triton X-100 (0.5%) in PBS. After incubation in blocking with 5% BSA in 0.05 % PBS-T for 60 minutes at RT, splenocytes were incubated overnight at 4°C with α-γH2AX (1/500, Cat # 20E3, rabbit monoclonal, CST, US) primary antibody. After washing with PBS followed by a 1h incubation with a goat anti-Rabbit Alexa-568-conjugated IgG (1/500, Invitrogen, Cat # A11011) at RT. After washing several times with PBS, the section was dried and mounted in DAPI-containing media. Images were captured with a Zeiss LSM 780 confocal microscope. The quantification of γH2AX foci intensity and number were performed using custom-trained Cellpose deep-learning model, and Random-Forest ML pixel-classifier.

#### Flow cytometry of spleen

All anti-mouse antibodies used were CD8-PE (Cat # 552094), CD4-FITC (Cat # 553650), CD3e-APC (Cat # 553066), and B220-APC-Cy7 (Cat # 552094), all purchased from BD Pharmingen. Spleen cells were isolated and red blood cells were removed using ACK lysis buffer. The stained cells were resuspended in FACS buffer (PBS with 0.5% BSA) to a final concentration of $10^7$ cells/mL. A 100 µL aliquot of the cell suspension was blocked with Fc blocker (anti-CD16/32) for 15 minutes at 4°C, then incubated with either single antibodies or an antibody cocktail for 30 minutes at 4°C. After incubation, cells were washed once with FACS buffer. Stained cell populations were analyzed using a BD LSRFortessa and FlowJo 10. Flow cytometry gating strategies are shown in Supplementary Fig. S3.

**Flow cytometry of immune subset of liver**

Freshly resected mouse liver tissue samples were stored in the MACS tissue storage solution (Miltenyi #130-100-008) at $4^0$ C not longer than 24 hours prior to dissociation. Liver tissues were mechanically dissociated using mouse liver dissociation Kit (Miltenyi #130-105-807) using gentleMACS™ dissociator (Miltenyi #130-093-926) following manufacturer's protocol. Single cell suspension dissociated from liver tissues were processed for multiparameter flow cytometric analysis. Cells were incubated with Live/Dead fixable Aqua dead cell stain kit (Invitrogen #L34957). Prior to antibody staining cells were blocked using Fc receptor blocking agent, mouse (Miltenyi Biotec #130-092-575), followed by incubation with antibodies for surface staining for 20 min at 4 °C. The antibodies used were CD4 (RM4-5 Biolegend #100542), CD8 (53-6.7 Biolegend #100733), CD3 (17A2 Biolegend #100237), CD11b (1D3/39 Biolegend #152408), CD11c (N418 Biolegend#117310), Ly6C (HK1.4 Biolegnd #128031), Ly6G (1A8 Biolegend #127618), MHC class II (REA813 Miltenyi #130-112-391),), CD45 (30-F11 Biolegend #103151), CD19 (1D3/CD19 Biolegend #152413). All flow cytometric analyses were performed using a MACSQuant Analyzer (Miltenyi Biotec) and data were analyzed using FlowJo software version 10.6.1. (FlowJo, LLC). Flow cytometry gating strategies are shown in Supplementary Fig. S4B.

**Interleukin-6 Measurements**

Prior to euthanasia, aliquots of whole blood were obtained from anesthetized mice for hematocrit determination and plasma preparation. IL-6 level was measured by BioTechne Luminex Mouse Discovery Assay according to the manufacturer's instructions.

**Complete blood count**

The mice were anesthetized using isoflurane anesthesia, then blood samples were collected through cardiac puncture before mice were euthanized. A total of 100 µl blood was collected and placed in the 0.5 ml EDTA-coated tube and shipped to MHL, NCI, Frederick for CBC.
